# Identification of a basal stem cell subpopulation in the prostate via functional, lineage tracing and single-cell RNA-seq analyses

**DOI:** 10.1101/601872

**Authors:** Xue Wang, Haibo Xu, Chaping Cheng, Zhongzhong Ji, Huifang Zhao, Yaru Sheng, Xiaoxia Li, Jinming Wang, Yu Shu, Yuman He, Liancheng Fan, Baijun Dong, Wei Xue, Chee Wai Chua, Dongdong Wu, Wei-Qiang Gao, Helen He Zhu

## Abstract

The basal cell compartment in many epithelial tissues such as the prostate, bladder, and mammary gland are generally believed to serve as an important pool of stem cells. However, basal cells are heterogenous and the stem cell subpopulation within basal cells is not well elucidated. Here we uncover that the core epithelial-to-mesenchymal transition (EMT) inducer Zeb is exclusively expressed in a prostate basal cell subpopulation based on both immunocytochemical and cell lineage tracing analysis. The Zeb1^+^ prostate epithelial cells are multipotent prostate basal stem cells (PBSCs) that can self-renew and generate functional prostatic glandular structures with all three epithelial cell types at the single-cell level. Genetic ablation studies reveal an indispensable role for Zeb1 in prostate basal cell development. Utilizing unbiased single cell transcriptomic analysis of over 9000 mouse prostate basal cells, we find that Zeb1^+^ basal cell subset shares gene expression signatures with both epithelial and mesenchymal cells and stands out uniquely among all the basal cell clusters. Moreover, Zeb1^+^ epithelial cells can be detected in mouse and clinical samples of prostate tumors. Identification of the PBSC and its transcriptome profile is crucial to advance our understanding of prostate development and tumorigenesis.

## Introduction

The prostatic epithelium, comprised of basal cells, secretory luminal cells and rare neuroendocrine cells, can serve as an excellent model to study stem cell biology due to its ability to regress and regenerate after repeated rounds of androgen deprivation and restoration in mice and rats ^1, 2^. Using sub-renal capsule tissue regeneration, lineage tracing, cell division mode analyses or organoid-forming assays, we and others have demonstrated the existence of prostate stem/progenitor cells in both basal and luminal prostate epithelia ^3–16^. However, none of the existing markers for basal stem cells (CD117, CD133, CD44, Trop2, CD49f, Sca1, etc.) ^8, 15, 16^ is exclusively found in the basal cell compartment. Despite the long-term postulation that basal cells contain more primitive prostate stem cells due to their resistance to castration, capability to differentiate into basal, luminal and neuroendocrine cell lineages of the prostate epithelium, and susceptibility to oncogenic transformation^6, 8, 15, 17–21^, the identity and nature of the prostate basal stem cells (PBSCs) within the heterogenous basal cell epithelia have not been elucidated.

In the present study, we identify a PBSC subpopulation that expresses Zeb1, an important EMT inducer ^22, 23^, through both *in vitro* and *in vivo* functional, lineage tracing and genetic ablation analyses. Furthermore, we integrated single-cell RNA sequencing with computational data analysis, a powerful approach that is lately developed to map cellular diversity of a given tissue, i.e., cortical neurons^24^ and cancer related immune cells^25^, etc., to examine the prostate basal cell compartment. Our single-cell transcriptomics data provide additional supporting evidence for the existence of Zeb1-expressing PBSCs and uncover their gene expression profile.

## Results

### Zeb1 is exclusively expressed in a prostate basal cell subpopulation and more frequently detected in the urethra-proximal region

EMT has been previously considered to occur only in early embryonic development or pathological conditions, including tumor metastasis and wound healing ^26^. Expression and function of core transcriptional factors such as Zeb1/2, Snai1/2, Twist1/2 which are required for EMT induction, have not been examined in normal prostate epithelia under physiological condition. Compared to prostate luminal cells, basal cells expressed low levels of epithelial-cell specific genes such as miRNA200 family^27^ and E-cadherin^3^ and exhibited a more mesenchymal-like phenotype. We therefore asked whether core EMT inducers are present in normal prostate basal cells. Immunofluorescent analysis of prostate sections showed that less than 1% of basal cells were positively labeled for Zeb1 immunostaining (Fig. 1a, b). Snai1 or Slug (Snai2) was expressed in 20% or 90% of p63^+^ basal cells, respectively, while Twist1/2 positive staining was not detected in the basal layer (Fig. 1a, b and Supplementary Table 1). Considering the interesting expression pattern of Zeb1 in the prostate epithelium and the previously reported role of Zeb1 in stemness acquisition and maintenance of cancer stem cells (CSC) characteristics^23, 28, 29^, we decided to investigate the biological relevance of Zeb1^+^ basal cells in the prostate.

**Fig. 1.**
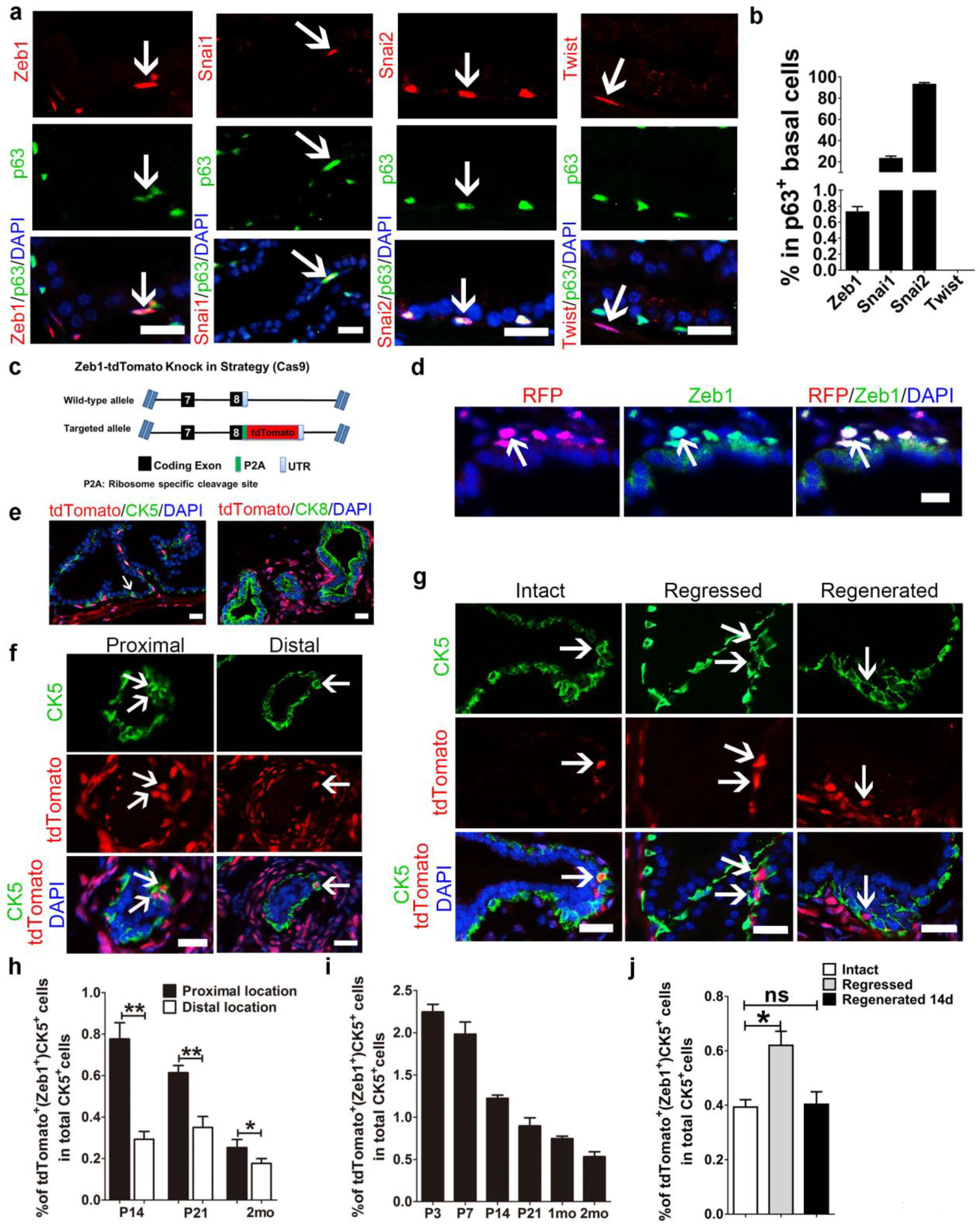
Mouse prostates contain a population of Zeb1^+^ basal cells that are more frequently detected in the urethra-proximal region. **a**, Co-immunostaining for Zeb1, Snai1, Snai2 (Slug), or Twist1/2 together with p63 on frozen sections of p21 wild-type mouse prostates. **b**, Quantification of the ratio of Zeb1^+^, Snai1^+^, Snai2^+^ or Twist1/2^+^ cells in p63^+^ prostatic basal epithelial cells. **c**, Illustration of the strategy to generate the Zeb1-tdTomato reporter mouse model. **d**, Zeb1 and tdTomato co-staining on prostate sections from Zeb1-tdTomato reporter mouse confirms that the td-Tomato labeling faithfully indicated the endogenous expression of Zeb1. **e**, Immunostaining of tdTomato and CK5 or CK8 on prostate frozen sections of 3-week-old wild type mice showing tdTomato expression is only found in prostate basal cells but not luminal cells. (Scale bars = 20μm) **f**, The tdTomato (Zeb1)^+^ prostate basal cells are more frequently found in the urethra-proximal region compared to the distal region. **g**, Section imaging of intact, regressed (21 days after castration) and regenerated (14 days after androgen replacement) prostate sections from Zeb1-tdTomato reporter mice. **h**, Quantification of tdTomato (Zeb1)^+^ prostate basal cells in urethra-proximal and distal regions at indicated developmental stages. **i**, Quantification of tdTomato (Zeb1)^+^ prostate basal cells at indicated time points of prostate development. **j**, Quantification of tdTomato (Zeb1)^+^ prostate basal cells at indicated time points of prostate regeneration. (In this figure, at least 20 fields per section of 3 sections each mouse prepared from 3 mouse prostates were analyzed. Data are analyzed by Student’s t-test and are presented as mean + s.e.m. All scale bars = 20μm.)

To better characterize and to more easily enrich the Zeb1^+^ prostate epithelial cells, we established a tdTomato knockin mouse model in which the coding sequence of tdTomato fluorescent reporter was linked to the last exon of *Zeb1* by a P2A element (Fig. 1c). Using immunofluorescent co-staining of Zeb1 and RFP on mouse prostate sections, we confirmed that the tdTomato labeling faithfully reflected the endogenous expression of Zeb1 (Fig. 1d). TdTomato positive cells were only found in prostate basal cells (marked by CK5 immunostaining) but not from luminal cell compartment (labeled by CK8 immunostaining) (Fig. 1e). The Zeb1^+^/tdTomato^+^ prostatic basal cells were more frequently detected in the urethra-proximal region relative to the distal region, the location where prostate stem cells were suggested to reside ^8, 30, 31^ (Fig. 1f, 1h and Supplementary Table 1). In addition, we found that the percentage of Zeb1^+^/tdTomato^+^ basal cells declined from 2.2% at postnatal day 3 (P3) to 1.0% at P21 as the prostate development proceeded, and decreased to 0.5% at adulthood (Fig. 1i and Supplementary Table 1). We then examined the dynamics of Zeb1^+^ basal cells during prostate regression and regeneration. As shown in Fig. 1g, 1j and Supplementary Table 1, the proportion of the Zeb1^+^CK5^+^ population became augmented in regressed prostates, and then decreased to the intact prostate level after regeneration, suggesting that they are more resistant to castration.

### Zeb1^+^ basal cells are enriched for multipotent prostate basal stem cells

To test the role of Zeb1^+^ basal cells in prostate development, we performed a prostate organoid-forming assay *in vitro* using flow cytometry sorted Zeb1^+^ and Zeb1^−^ basal cells (Fig. 2a, b). While few and small organoids were produced from Zeb1^−^ basal cells, significantly larger and more organoids were generated from Zeb1^+^ basal cells (Fig. 2c, 2e and Supplementary Table 1). Immunostaining analysis of frozen sections of organoids formed from sorted Zeb1^+^ basal cells showed generation of both basal (CK5, CK14 or p63 positive) and luminal (CK8 or AR positive) cells (Fig. 2d). Moreover, Zeb1^+^ basal cells possessed a serial organoid forming capacity, indicating a self-renewing characteristic (Fig. 2e and Supplementary Table 1). We then evaluated the potential of Zeb1^+^ basal cells to generate prostates *in vivo*, a gold standard to assess the stem cell phenotype ^8, 32, 33^. Using an *in vivo* kidney capsule transplantation assay, we grafted 1000 Zeb1^+^ or Zeb1^−^ mouse prostate basal cells in combination with rat embryonic urogenital sinus mesenchymal (UGM) stromal cells under the renal capsule of host athymic nude mice. Compared to opaque and small grafts derived from Zeb1^−^ basal cells, Zeb1^+^ basal cells generated semi-translucent and large grafts (Supplementary Fig. 1a-c). Histological analysis indicated that Zeb1^+^ grafts underwent ductal morphogenesis with differentiation of basal (CK5^+^), luminal (CK8^+^) and rare neuroendocrine (Syp^+^) cell lineages (Supplementary Fig. 1d, 1f). Pbsn expression further confirmed differentiation of functional secretory luminal cells (Supplementary Fig. 1f). In addition, Zeb1^+^ grafts expressed the mouse β-integrin, which validated their mouse origin and indicated that they were not produced from the contamination of rat epithelial cells from the preparation of rat UGM cells (Supplementary Fig. 1f). We further verified that the generated prostate tissues were originated from Zeb1^+^ donor cell grafts by positive staining of an MHC class I haplotype H-2k^b^ protein that is specifically expressed in C57BL/6 but not host athymic mouse cells (Supplementary Fig. 1f). These data suggested that Zeb1^+^ basal cells are multipotent prostate stem cells and possessed the capacity of generating functional prostates.

**Fig. 2.**
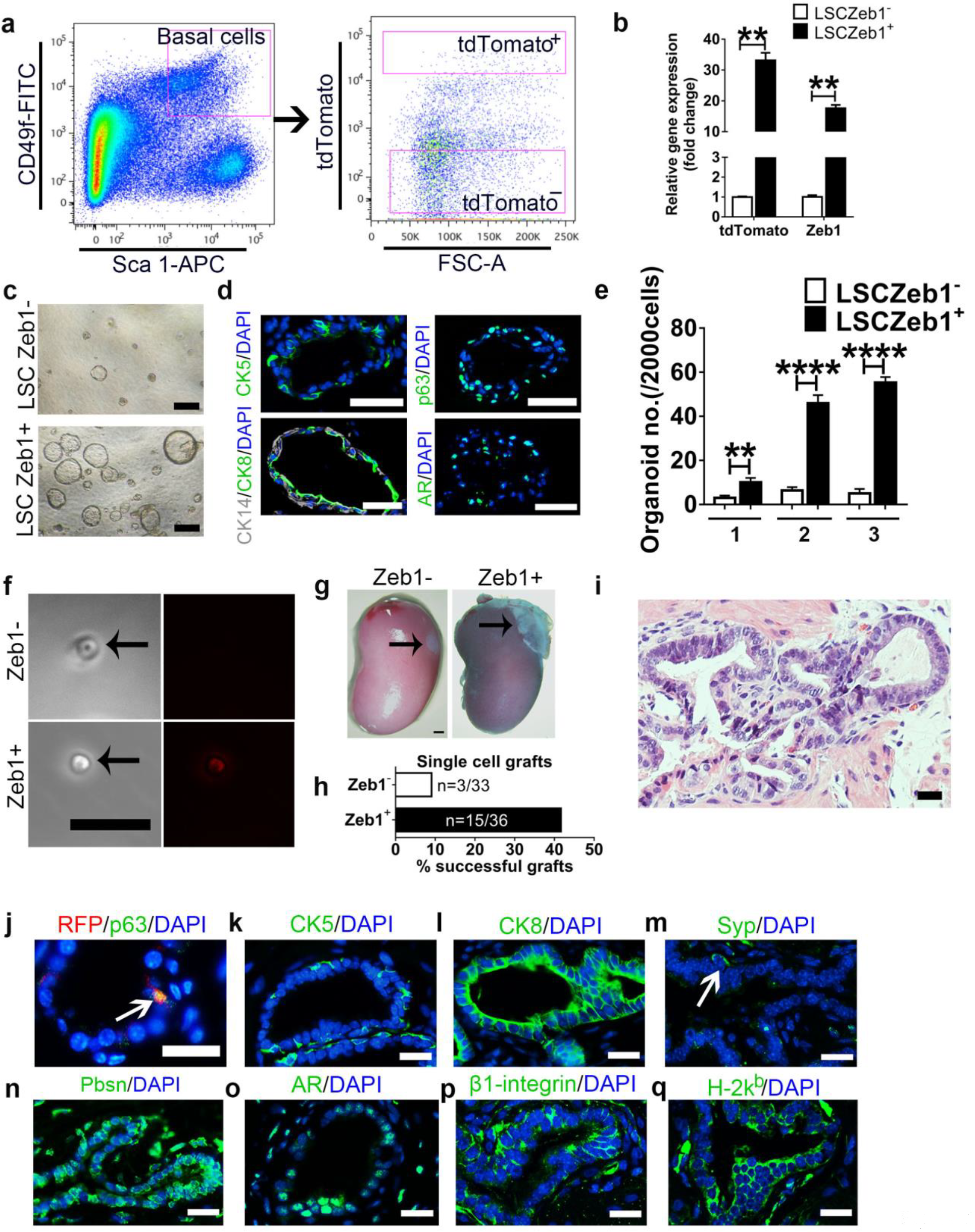
Zeb1^+^ basal cells are enriched for prostate stem cells. **a, b**, FACS and qRT-PCR quantification of Zeb1expression in Lineage^−^Sca-1^+^CD49f^hi^ tdTomato^+^ and Lineage^−^Sca-1^+^CD49f^hi^ tdTomato^−^ prostate cells. RNA expression levels were normalized to Lineage^−^Sca-1^+^CD49f^hi^ tdTomato^−^ prostate basal cells. (n=5) **c-e**, *In vitro* serial organoid-forming assay showing more and larger organoids were produced from Zeb1^+^ basal cells compared to Zeb1^−^ basal cells. (Scale bars = 200μm for bright-field images and 50μm for the immunofluorescent images. The number 1,2,3 stands for the first, secondary and tertiary passaging.) **f**, Phase contrast and fluorescent images of the single viable Lineage^−^Sca-1^+^CD49f^hi^ Zeb1^+^ or Lineage^−^Sca-1^+^CD49f^hi^Zeb1^−^ prostate cell used for single cell renal capsule transplantation experiments *in vivo*. **g**, Prostate tissue generated from single Lineage^−^Sca-1^+^CD49f^hi^ Zeb1^+^ prostate cell at 2 months after renal capsule implantation. **h**, Quantification of prostate tissue generation incidence from single cell transplants. **i**, H&E staining of single cell implants showing differentiated prostate epithelial tubules. **j-o**, Staining of RFP, p63 (**j**, green), CK5 (**k**), CK8 (**l**), Synaptosphysin (Syp) (**m**), Pbsn (**n**), and Androgen receptor (AR) (**o**) on sections of single cell implants. (The white arrow points to a Syp^+^ cell. Scale bars=20 μm.) **p, q**, Staining of mouse-specific β1-integrin (**p**) and C57BL/6 donor-specific H-2k^b^ (**q**) on sections of single cell implants. (Scale bars=20 μm.) (data are analyzed by Student’s t-test and are presented as mean + s.e.m.)

### Single Zeb1^+^ basal cell can generate functional prostate *in vivo*

We then wondered whether a single Zeb1^+^ basal cell would generate a prostate *in vivo*. Single viable Zeb1^+^ or Zeb1^−^ basal cell was FACS sorted into individual well of a 96-well plate, mixed with rat UGM stromal cells and transplanted into the renal capsules of nude mice. Each well was examined under the microscope to confirm the presence of a tdTomato^+^ or tdTomato^−^ single cell (Fig. 2f). Fifteen prostates with well-differentiated prostate epithelial tubules containing all three basal, luminal and neuroendocrine cell lineages were generated from 36 single Zeb1^+^ basal cell transplants (Fig. 2g-q). In contrast, only 3 prostates were produced from 33 single Zeb1^−^ basal cell xenografts (Fig. 2g-q). Furthermore, Zeb1^+^Epcam^+^ cells harvested from the first generation of xenografted prostate tissues were able to form functional prostates in the secondary and tertiary sub-renal capsule transplantations (Supplementary Fig. 2). Collectively, these data provided additional direct evidences that Zeb1^+^ basal cells represented a multipotent prostate stem cell population.

### *In vivo* lineage tracing supports that Zeb1^+^ basal cells give rise to basal, luminal and neuroendocrine progeny

To further determine the role of Zeb1^+^ basal cells in prostate development *in vivo*, we generated a *Zeb1-CreERT2* mouse line and crossed it with *Rosa-CAG-LSL-tdTomato* mice to trace Zeb1 ^+^ basal cells during the postnatal prostate development (Fig. 3a, b). Tamoxifen administration to postnatal day 3 *Zeb1-CreERT2/tdTomato* mice induced expression of tdTomato in 10% of Zeb1^+^ cells (Fig. 3c). Consistent with our immunostaining results and findings from the Zeb1-tdTomato knockin mouse model (Fig. 1), about 0.33% of CK5^+^ basal cells were labeled by tdTomato but none of the CK8^+^ luminal cells nor Syp^+^ neuroendocrine cells were labeled at 2 days after induction (Fig. 3d-f, 3i and Supplementary Table 1). Interestingly, 12 days after the tamoxifen administration, we found clusters of CK5^+^CK8^−^tdTomato^+^ cells located in the outer basal cell layer as well as CK5^−^ CK8^+^tdTomato^+^ cells in the inner luminal compartment (Fig. 3g, 3i and Supplementary Table 1). Quantitative analysis exhibited that labeled basal cells expanded more than 16 folds and tdTomato expressing luminal cells increased from zero to around 4% in 12 days. Meanwhile, we also observed that about 3% of Syp^+^ neuroendocrine cells were marked with tdTomato at 12 days post induction (Fig. 3h). Those results further substantiated the notion that Zeb1^+^basal cells were able to generate basal, luminal and neuroendocrine cell lineages.

**Fig. 3.**
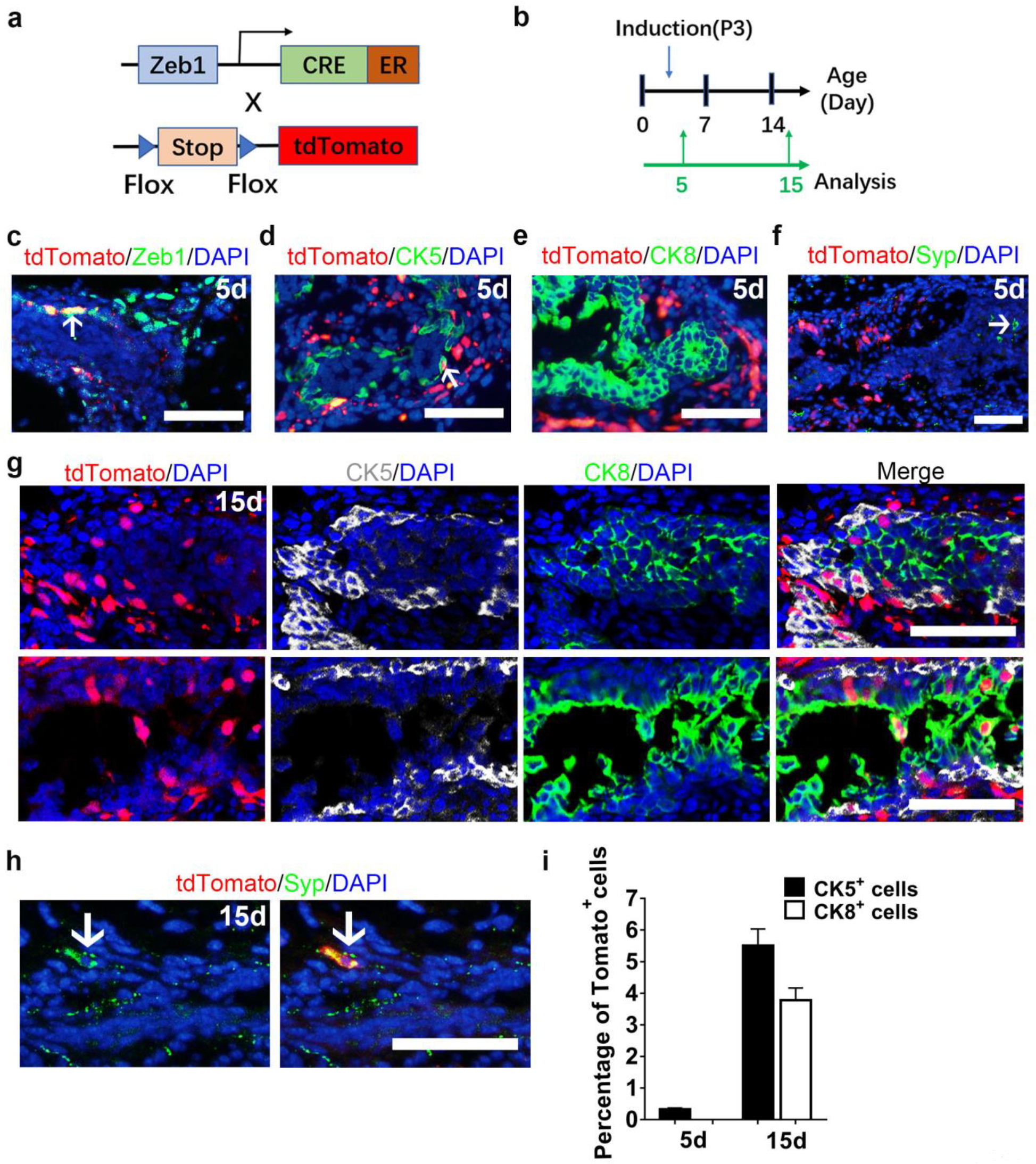
Lineage tracing supports that Zeb1^+^ basal cells give rise to basal, luminal and neuroendocrine cells. **a**, Strategy to target tdTomato expression in Zeb1 expressing basal cells *in vivo*. **b**, Illustration of protocols to trace the fate of Zeb1^+^ basal cells during prostate postnatal development. **c**, Zeb1 and tdTomato co-staining of prostate sections from *Zeb1-CreERT2/tdTomato* mice shows that tdTomato marks Zeb1 expressing cells. **d-f**, tdTomato expression is found in CK5^+^ basal cells, but not in CK8^+^ luminal cells and Syp^+^ neuroendocrine cells from *Zeb1-CreERT2/tdTomato* mouse prostates at 2 days after tamoxifen administration. **g**, Clusters of CK5^+^CK8^−^tdTomato^+^ cells can be detected in the outer basal cell layer and CK5^−^CK8^+^tdTomato^+^ cells can be found in the inner luminal compartment at 12 days after tamoxifen induction. **h**, Syp^+^ neuroendocrine cells are marked with tdTomato at 12 days post induction. **i**, Percentage of tdTomato^+^ cells in CK5^+^ and CK8^+^ cells at 2 days and 12 days after tamoxifen administration to postnatal day 3 mice. (In this figure, at least 20 sections each mouse prepared from 3 mouse prostates were analyzed. At least 60 fields for each immunostaining experiment were collected for analysis. Data are analyzed by Student’s t-test and are presented as mean + s.e.m. All scale bars = 50μm.)

### Zeb1 is required for development of prostatic basal cells

We then asked whether Zeb1 was functionally required for normal prostate development. To that end, we first utilized the clustered regularly interspaced short palindromic repeats (CRISPR)/CRISPR-associated protein 9 (Cas9) system to achieve *Zeb1* gene knockout in prostate organoids *in vitro* (Fig. 4a). Efficient *Zeb1* deletion in mouse primary prostate epithelial cells was validated by immunoblotting (Fig. 4b). Zeb1 sgRNA transfected prostate organoids contains CK8^+^ luminal cells, however, Zeb1 knockout organoids displayed a marked decrease of p63 expressing basal cells compared to organoids formed from mock-transfected prostate epithelial cells (Fig. 4c, d and Supplementary Table 1). We next investigated the role of Zeb1 in prostate development *in vivo*. Due to the fact that Zeb1 knockout mice died shortly after birth^34^, UGS from E16 Zeb1^−/−^ embryos were dissected and transplanted beneath the renal capsule of male athymic mice to allow us to assess prostate epithelial development with Zeb1 deletion (Fig. 4e). Although Zeb1^−/−^ UGS grafts developed into prostate and contains prostatic ductal structures, Zeb1 null grafts were smaller (Fig. 4e, f). Detailed histological examination revealed that approximately 50% of the Zeb1^−/−^ UGS derived prostate epithelia virtually displayed all luminal cells that are positive of CK8 and AR without basal cells. In the other prostate epithelia, basal cell numbers were greatly reduced (Fig. 4f-h and Supplementary Table 1). Further immunofluorescent co-staining with Zeb1 and p63 antibodies affirmed Zeb1 deletion and marked decrease of basal cells in Zeb1^−/−^ UGS grafts (Fig. 4g, h and Supplementary Table 1). These findings were further supported with additional staining of other basal cell markers CK5 and CK14 (Fig. 4g, h and Supplementary Table 1). Syp-expressing neuroendocrine cells could still be found in Zeb1^−/−^ UGS derived prostate epithelium (Fig. 4g). Collectively, our *in vitro* and *in vivo* data highlight an indispensable role for Zeb1 in prostate basal cell development.

**Fig. 4.**
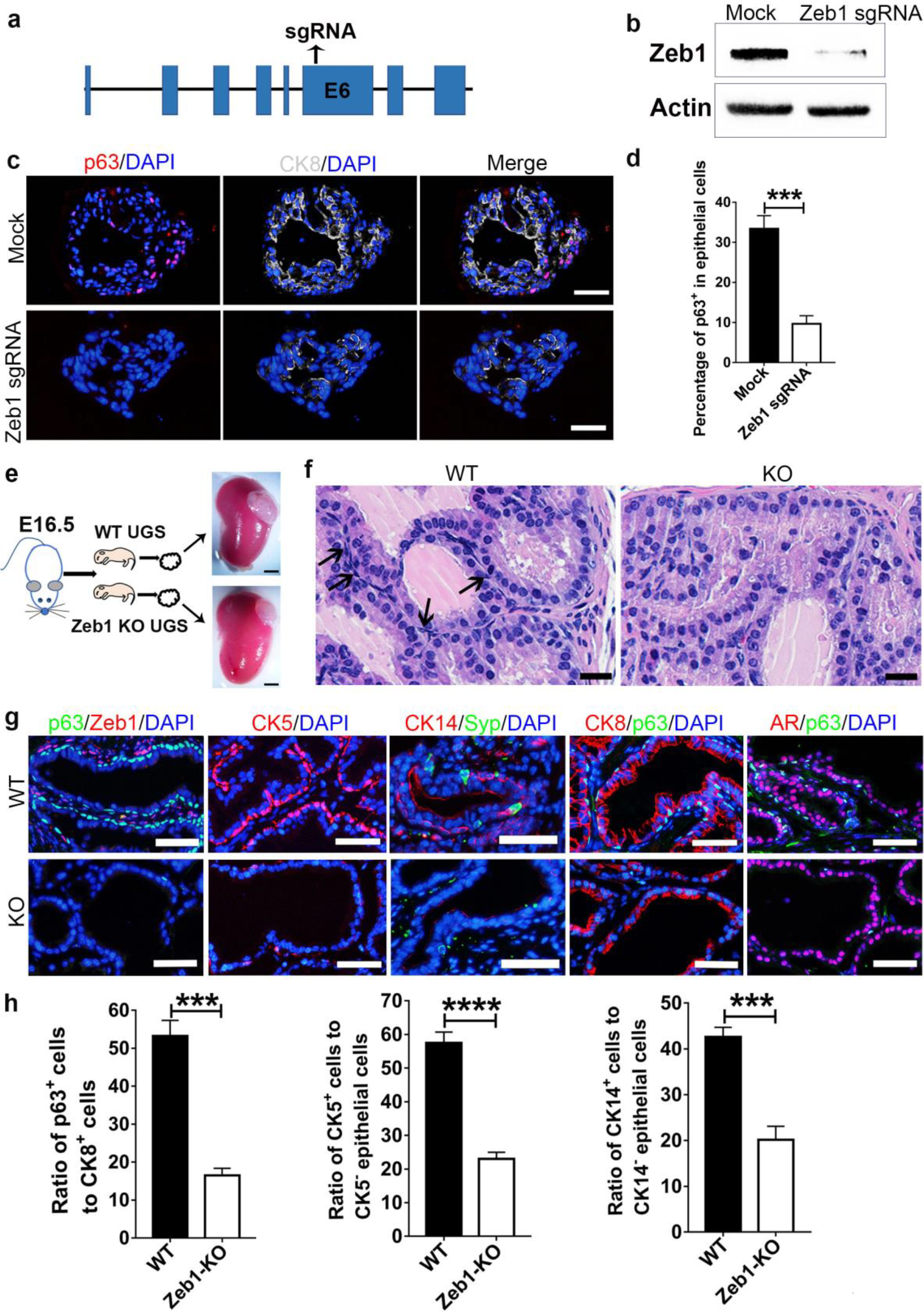
Knockout of Zeb1 severely suppresses the basal cell development. **a**, Illustration of sgRNA sequence for CRISPR/Cas9 system mediated Zeb1 knockout. **b**, Immunoblotting confirms efficient Zeb1 ablation sgRNA transfected primary prostate epithelial cells. **c, d**, Immunostaining of frozen sections from Zeb1 knockout organoids shows a severely impaired differentiation of basal cells. (Scale bars =50μm. Experiments were repeated for 3 times. Data are analyzed by Student’s t-test and are presented as mean + s.e.m.) **e**, UGS from E16 wild-type or Zeb1^−/−^ embryos are dissected and transplanted beneath the renal capsule of male athymic mice. Transplants are harvested at 20 days later. (Scale bars =200μm) **f**, H&E staining of UGS implants showing differentiated prostate epithelial tubules but a remarkably decrease in basal cell number in Zeb1^−/−^ UGS grafts. (Scale bars =20μm) **g**, Staining of Zeb1, p63, CK5, CK14, Syp, CK8 and AR on sections of wild-type and Zeb1 knockout UGS implants. (Scale bars =50μm) **h**, Quantification of basal cell development in Zeb1^−/−^ UGS grafts. (At least 60 fields from 20 sections for each immunostaining experiment were analyzed. Data are analyzed by Student’s t-test and are presented as mean + s.e.m.)

### Unbiased single cell transcriptomic analysis of prostate basal cells identifies a unique Zeb1^+^ cell subset which expresses both epithelial and mesenchymal gene signatures

To provide additional evidence for the existence of a Zeb1^+^ basal population, we performed unbiased single cell transcriptomic analysis of prostate basal cells. Using the 10x Genomics single cell system followed by Illumina sequencing, prostate basal cells (lineage^−^Sca1^+^CD49f^hi^) ^15^ were purified from mouse prostates via FACS for single-cell RNA sequencing. Sorted basal cells were confirmed by q-RT-PCR and immunostaining for expression of basal cell markers (Supplementary Fig. 3a-c). The flow chart for single-cell RNA sequencing data analysis was included in Supplementary Fig. 3d. To visualize cellular heterogeneity, 9833 single-cell transcriptome data were subjected to unsupervised Seurat clustering and projected onto two dimensions by t-distributed stochastic neighbor embedding (t-SNE). After removal of low-quality and contaminated non-epithelial cells (Supplementary Fig. 3e, f and 4), this led to a total of 9 prostate basal cell clusters (Cluster 1-9 or termed as C1-C9) (Fig. 5a). Importantly, we further found that while all other basal cell clusters displayed high levels of epithelial genes, the cell cluster C7, expressed both epithelial markers *Epcam*, *Cldn1*, *Ocln* and stromal cell markers including *Vim*, *Fn1*, and *Acta2*, stood out uniquely among all the cell clusters (Fig. 5a, b and Supplementary Table 2). Detailed analysis of highly variable genes in C7 showed that Zeb*1* was exclusively expressed in C7 (Fig. 5a, b and Supplementary Table 2). Other EMT transcriptional factors including *Zeb2*, *Snai1, Prrx1* and *Prrx2* were also highly transcribed in C7 (Fig. 5a, b and Supplementary Table 2). This EMT-like cell cluster can also be identified through the Monocle cell clustering method (Supplementary Fig. 5a, b). Interestingly, when we used statistical and computational frameworks to delineate the differentiation states of the 9 cell clusters, both monocle and diffusion map scripts indicated that C7 was at one of the tips in the lineage tree (Supplementary Fig. 5). Collectively, the unbiased examination of single-cell gene expression profiles of prostate basal cells further substantiates a unique Zeb1 expressing basal cell subpopulation with the mesenchymal gene signature.

**Fig. 5.**
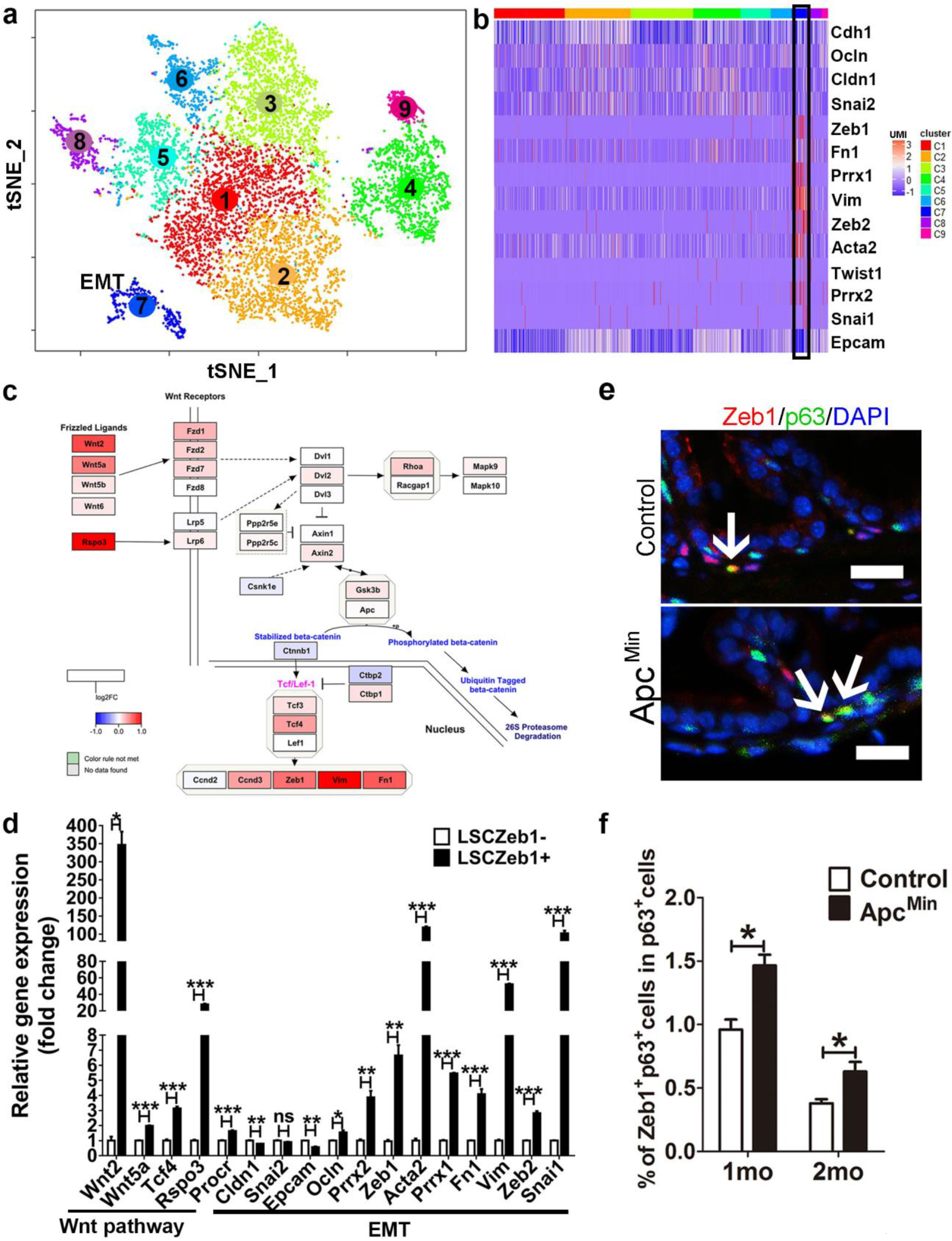
Single-cell RNA-seq data reveal that a Zeb1^+^ basal cell subset which shares gene transcriptional signatures with both epithelial and mesenchymal cells meanwhile preferentially expresses molecules in the Wnt signaling pathway and is expanded in APC^min^ mice. **a**, A Seurat package and the first 12 principal components were applied to generate 9 different and stable clusters based on differential expressing genes among 9278 mouse prostate basal cells. C7 is marked with EMT due to its unique EMT expression profile. **b**, A heatmap showing the scaled expression profile for EMT related genes. (The black-lined rectangle highlights cluster 7). **c**, Color-coded expression levels of Wnt signaling pathway genes in the Zeb1 expressing Cluster 7 were presented as were obtained according to log2 fold change (log2FC) between cluster 7 and the rest prostate basal cell clusters. The pathway drawing was modulated from the schematic for the mouse Wnt pathway at the pathviso website. **d**, qRT-PCR quantification of the mRNA expression of key components of the Wnt pathway and epithelial and stromal markers as well as EMT inducing transcriptional factors in Lineage^−^Sca-1^+^CD49f^hi^ Zeb1^+^ and Lineage^−^Sca-1^+^CD49f^hi^ Zeb1^−^ prostate cells. (n=3, data are analyzed by Student’s t-test and are presented as mean + s.e.m.) **e, f**, Co-immunostaining of Zeb1 and p63 on prostate sections reveals a significant increase of Zeb1^+^ basal cell number in APC^min^ mice compared to control animals. (Scale bars =20μm)

### Wnt signaling pathway is enriched in Zeb1^+^ basal cells

As an effort to uncover signaling pathways that are enriched in the cell cluster C7, we applied gene ontology (GO) enrichment analysis to the single-cell RNA-seq data. Coincidently, we found that the Wnt signaling pathway, which we have reported recently to promote self-renewal of prostate cancer stem cells ^35^, was one of the top hits preferentially presented in the cluster C7 (Fig. 5c and Supplementary Table 3). The Wnt signaling pathway was demonstrated previously to play an essential positive role in inducing EMT ^36^. Our RT-PCR experiments confirmed a significant upregulation of key Wnt signaling components, EMT transcriptional factor and mesenchymal markers, and downregulation of epithelial makers in Zeb1^+^ basal cells (Fig. 5d). It was recently reported that Procr, a Wnt pathway target, marks a novel multipotent mammary stem cell population ^37^. Consistently, we also found a moderate (1.5 fold) but significant upregulation of Procr in Zeb1^+^ prostate basal cells (Fig. 5d). Further flow cytometry examination revealed that Zeb1^+^ cells represented only a small subpopulation (8%) of Procr expressing prostate epithelial cells (Supplementary Fig. 6), suggesting a more enrichment of PBSCs in Zeb1^+^ than Procr^+^ prostate epithelial cells. Furthermore, examination of prostate sections from APC^min^ mice with the aberrantly activated Wnt pathway revealed a significant increase of Zeb1^+^ basal cells (Fig. 5e, f and Supplementary Table 1), suggesting a positive role of the Wnt signaling in the expansion of Zeb1^+^ basal cells.

**Fig. 6.**
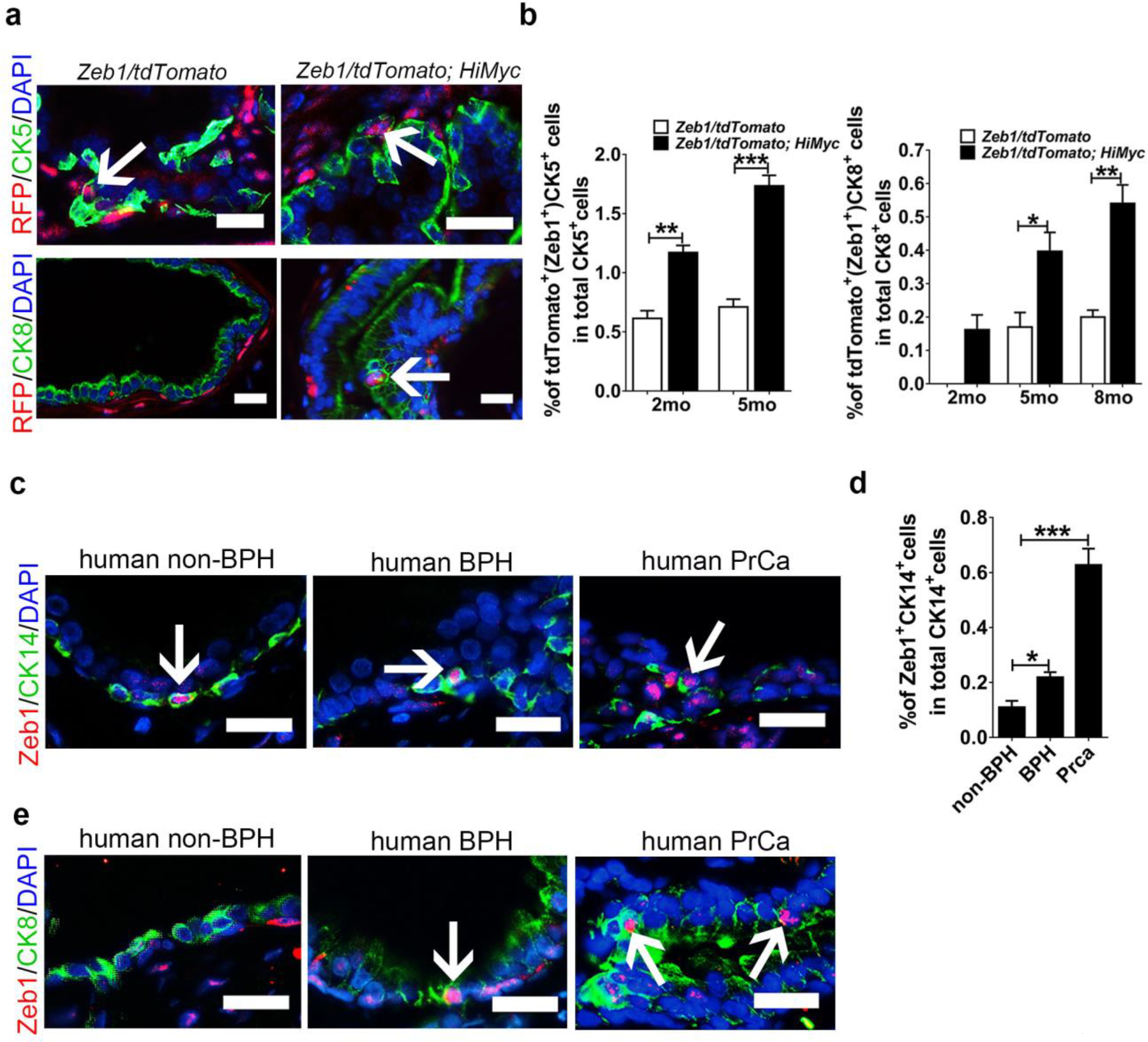
Zeb1^+^ epithelial cells are found in the basal layer of both mouse and human prostates and are detectable in both basal and luminal epithelium from mouse or human hyperplastic prostate tissues. **a**, Images of tdTomato and CK5 or CK8 staining on prostate sections from *Zeb1/tdTomato or Zeb1/tdTomato; HiMyc* mice. **b**, Quantifications of tdTomato^+^ basal or tdTomato^+^ luminal cells in prostates from *Zeb1/tdTomato or Zeb1/tdTomato; HiMyc* mice. (In panel **a-d**, at least 20 fields per section of 3 sections each mouse prepared from 3 mouse prostates were analyzed. Data are analyzed by Student’s t-test and are presented as mean + s.e.m. All scale bars = 20μm.) **c, d**, Images and quantifications of Zeb1^+^ basal epithelial cells in human non-BPH, BPH or prostate cancer specimens. **e**, Images of Zeb1 and CK8 co-immunostaining on sections from human non-BPH, BPH or prostate cancer specimens (In panel **c-e**, non-BPH (n=3), BPH (n=7). For cancer samples, Gleason score=3 + 3 or 3+ 4, n=4. Scale bars=25μm).

### Zeb1-expressing cells can be detected in prostate epithelia of *Hi-Myc* mice and human prostate samples

We next asked the influence of oncogenic transformation on the Zeb1 expression profile in the prostate epithelium. For that purpose, we crossed the Zeb1/tdTomato mice with the *Hi-Myc* prostate cancer mouse model. Immunofluorescent co-staining of Zeb1 or tdTomato together with basal or luminal cell markers revealed a 2-fold increase of Zeb1^+^ basal cells in *Zeb1/tdTomato; HiMyc* prostates (1.25% versus 0.73% in control mice at 3-month old, or 1.71% versus 0.71% in control mice at 5-month old) (Fig. 6a, b and Supplementary Table 1). Of note, we detected a small number of Zeb1^+^CK8^+^ luminal cells in *HiMyc* mice, which stood in contrast to the exclusive expression of Zeb1 in the basal cell compartment from wild type prostates (Fig. 6a, b and Supplementary Table 1). Similarly, positive immunostaining of Zeb1 protein could be found in a small percent of CK14^+^ basal cells in human prostate tissues (non-benign prostate hyperplasia (non-BPH) (n=3) and BPH specimens (n=7)) (Fig. 6c, d and Supplementary Table 1). During the examination of many sections of human prostate cancer samples, we infrequently found cancer tissues (Gleason score: 3+3 or 3+4, n=4) with remaining CK14^+^ basal cells, in which Zeb1 positive staining could be found (Fig. 6c, d and Supplementary Table 1). Intriguingly, Zeb1^+^ CK8^+^ luminal cells were barely detected in non-BPH but could be observed occasionally in BPH specimens and more frequently in prostate cancer specimens (Fig. 6e). Together, Zeb1 expressing epithelial cells can be detected in both basal and luminal layers of prostates from *Hi-Myc* mice and in human prostate samples.

## Discussion

The prostate basal cell compartment is suggested to contain stem/progenitor cells ^6, 8, 15, 21^, but the stem cell subpopulation within prostate basal cells and its transcriptional profile remain largely unknown. We here discover the exclusive existence of Zeb1-positive prostate epithelial cells in a small population of basal compartment but not in the luminal layer. Zeb1^+^ basal cells are more frequently found in the stem cell enriched urethra-proximal region. Importantly, utilizing functional methods and *in vivo* lineage tracing, we prove that the Zeb1^+^ prostate basal cells can self-renew and possess multipotency to generate all three prostatic epithelial cell lineages *in vivo*. Furthermore, *in vitro* and *in vivo* genetic ablation experiments showed a requirement of Zeb1 in basal cell differentiation. The unique Zeb1 expressing prostate basal cell cluster with mesenchymal cell features was further supported by single-cell RNA sequencing data. Our study underscores the importance of Zeb1 as a single marker and an essential regulator for PBSCs.

EMT has been demonstrated to be utilized by tumor initiating cells to acquire stem cell attributes ^23^. We find here that Zeb1 marks a population of PBSCs in prostates with mesenchymal features and higher expression levels of EMT inducers such as Snai1, Zeb2, Prrx1 and Prrx2. Support for our data comes from recent studies in mammary glands that multipotent Procr^+^ mammary stem cells display EMT characteristics and that EMT is required for the tissue-reconstitution activity of mammary basal cells ^37–40^. Together, those observations demonstrate that like what was shown for tumor initiating cells, EMT is also closely associated with stemness in normal prostate and mammary stem cells. However, there is also tissue specific difference between stem cells in prostate versus mammary tissues. Firstly, we find that although almost all Zeb1^+^ prostate basal cells express Procr, Procr is present in a much larger population of prostate epithelial cells containing both basal and luminal cells (Supplementary Fig. 6 and data not shown), suggesting that Zeb1 appears to be a better maker for basal stem cells in prostates than Procr. Secondly, Zeb1 and Snai1 are expressed in prostate basal cells but not in mammary epithelial cells ^38–40^, indicating distinct EMT inducing transcriptional factors or EMT programs are involved in respective stem cells.

The serial organoid forming and transplantation assays together with lineage tracing experiments of Zeb1^+^ basal cells in the current study demonstrate that Zeb1^+^ basal cells can self-renew and give rise to basal, luminal and neuroendocrine cell lineages. Loss of function experiments performed on prostate organoids and UGS transplants show that Zeb1 knockout results in a remarkably decrease in the basal cell compartment, but luminal and neuroendocrine cell differentiation are largely unaffected. Consistently, previous studies on renal capsule transplants from p63^−/−^ UGS exhibit development of luminal and neuroendocrine cells independent of basal cells ^41^. In addition, lineage tracing experiments on CK14, CK8, CK5 and NKX3.1 cells support the existence of self-sustained luminal cell progenitors ^5, 6, 9, 10^. Therefore, both multipotent Zeb1^+^ basal stem cells and stem/progenitor cells from the luminal compartment contribute to prostate development.

Intriguingly, we find that different from the exclusive basal expression of Zeb1 in wild-type prostate, Zeb1^+^ epithelial cells can be detected in the luminal compartment in Hi-myc mice or in human prostate hyperplastic or tumor samples. There are two possible explanations for the appearance of Zeb1^+^ luminal cells: It could be derived from transformed Zeb1^+^ basal cells; or alternatively, the transformed luminal cells may acquire Zeb1 expression. Clarification of these two possibilities needs further investigations. Nevertheless, the appearance of Zeb1^+^ luminal cells at early stages of prostate tumorigenesis is of special interest, because most clinical prostate cancers display a dominant luminal phenotype by loss of basal cell phenotypes, and Zeb1 has been well shown in the acquisition of cancer stem cell properties ^28, 42^. This Zeb1^+^ luminal cell population might represent prostate tumor initiating cells, which can be also related to two different cellular origins for prostate cancers: basal versus luminal cells.

To our knowledge, this work is the first report to decipher the heterogeneity within the prostate basal epithelium using unbiased single-cell transcriptomics. Importantly, along with the single-cell RNA-seq data, our immunocytochemical staining, functional experiments, and cell lineage tracing analysis together support that Zeb1 marks a multipotent prostate basal stem cells and Zeb1 itself is required for prostate basal cell development. The identification of the novel Zeb1^+^ prostate stem cell, its transcriptome profile and its expression pattern during prostate tumorigenesis could have important implications for our understanding of tissue development and regeneration and for identification of potential cellular origin for cancer.

## Supporting information

Supplemental info.(including supplemental figures and figure legends)

## Acknowledgements

The study is supported by funds to by funds to W-Q Gao from the National Key R&D Program of China (2017YFA0102900), the National Natural Science Foundation of China (NSFC, 81630073 and 81872406), the Science and Technology Commission of Shanghai Municipality (16JC1405700), the Education Commission of Shanghai Municipality (for the High Peak IV subject on Stem Cells and Translational Medicine Research) and the KC Wong foundation, and by funds to H.H. Zhu from the NSFC (81772743), Shanghai Municipal Education Commission— Gaofeng Clinical Medicine Grant Support (20181706), Shanghai Rising-Star Program (17QA1402100), the Shanghai Youth Talent Support Program, School of Medicine, Shanghai Jiao Tong University (Excellent Youth Scholar Initiation Grant 16XJ11003).

## Author contributions

H.H.Z. and W.Q.G. conceived the study; X.W. performed the experiments. H. X. and D. W. conducted bioinformatic analyses; Z. J. and J. W. assisted in data interpretation; C. C. and Y. S. helped in cell culture experiments; H.Z. helped microscopic imaging; X. L. performed mouse genotyping and breeding; Y. S. helped organoid culture; Y. H. assisted in animal experiments. L.F., B.D., and W.X. collected human prostate samples and assisted in histological analysis of human sample; H.H.Z., W.Q.G. and X.W. interpreted the data and wrote the manuscript.

## Methods

### Animals

Pregnant SD rats (E17) and athymic nu/nu male mice (7 weeks old) were purchased from Shanghai Slac laboratory animal company. *Zeb1/tdTomato* reporter mice were generated by the CRISPR-Cas9 method at Model Animal Research Center of Nanjing University. The sgRNA sequence we used is GAGGTTGGAGCTGCACAGCAGG. Exon 8 of Zeb1 followed by a P2A and tdTomato coding sequence, which was flanked by approximately 1.5kb upstream and 1.5kb downstream sequence of Zeb1 Exon 8, was cloned to a donor vector. Plasmid containing sgRNA and Cas9 expressing elements and the donor vector were injected to fertilized C57/B6 mouse eggs. F0 generation of mice were genotyped by sequencing and PCR. Positive F0 mice were used to backcross with C57/B6 mice to produce F1 generation of knockin mice. PCR Primers for *Zeb1/tdTomato* mouse genotyping were provided in Supplementary Table 4. *Zeb1-CreERT2* mice were generated by a homologous recombination method at the Cyagen Biosciences Inc. The TAG stop codon of the last exon of Zeb1 was replaced with the “2A-CreERT2” cassette. Homology arms of the targeting vector were generated by PCR using BAC clone RP23-51G9 or RP23-207F18 from the C57BL/6J library as template. The targeting vector contained the Neo cassette flanked by SDA (self-deletion anchor) sites and DTA for negative selection. C57BL/6 ES cells was used for gene targeting. F0 generation of mice were validated by sequencing and PCR. PCR Primers for genotyping of *Zeb1-CreERT2* mice were provided in Supplementary Table 4. *Zeb1* knockout with deletion of exon 1 mice was purchased from RIKEN BioResource Research Center. The *Rosa-CAG-LSL-tdTomato* mice were purchase from the Jackson laboratory. *Hi-Myc* mice were introduced from the National Cancer Institute (NCI:01XF5). *Apc*^*min*^ mice were introduced from the Nanjing Biomedical Research Institute of Nanjing University (T001457). All animal experiments were conducted according to the ethical regulations of Ren Ji Hospital. Animal experiment protocol were approved by the Ren Ji Hospital Laboratory Animal Use and Care Committee.

### Prostate single cell preparation

Mouse prostates were harvested, minced, then digested with pre-warmed 1X collagenase/hyaluronidase solution for 3hr at 37℃, washed with PBS buffer once and placed in 0.25% Trypsin/EDTA for 6 min at 37℃. FBS supplemented with 4% FBS was added to quench trypsin reaction followed by centrifuging at 350g for 5min. Then, the cell pellet was resuspended with pre-warmed Dispase/DNase I solution to thoroughly dissociate cells. Single cell suspension was obtained from passing through a 40µm cell strainer.

### Flow cytometry

Mouse prostate single cell suspension was blocked with the Fc blocker (CD16/32 antibody from eBioscience) for 40min at room temperature. Staining antibodies were diluted in 4% FBS buffer containing Y-27632 ROCK inhibitor and applied to the prostate single cell suspension for 40min at room temperature. Flow cytometry analysis or sorting was conducted using BD Accuri C6 or FACSAria III flow cytometer. Antibodies used in the study are listed in Supplementary Table 5. Propidium Iodide (PI, Invitrogen) was added to sample before sorting to gate viable cells.

### Single cell RNA-seq and data analysis

Lineage^−^Sca-1^+^CD49f^hi^PI^−^ prostate cells from postnatal day 14 mice were sorted and counted manually before processing to the single cell RNA-seq library preparation. Libraries were constructed following the instruction of the Chromium single cell 3’ solution (10x Genomics). The libraries were sequenced on Illumina Hiseq X Ten platform. 10x Genomics workflow and Cell Ranger Single Cell Software was used to process raw sequencing data and to align reads to the mm10 mouse reference genome. Raw gene expression matrix produced by 10x Genomics workflow were first processed by the Seurat package. Cells with less than 5000 unique molecular identifiers (UMIs) or less than 1400 genes detected or more than 5% UMI mapped to mitochondria genes were removed (Supplementary Fig. 3e, f). This led to 9833 high-quality single-cell RNA-seq data with an average gene detected to be around 2703 (Supplementary Fig. 3f). 1606 variable genes were selected based on their expression and dispersion (expression cutoff=0.0125 and dispersion cutoff=0.5). The first 12 principal components were used for clustering analysis (resolution=0.5) and t-SNE projection. This identified a total of 10 clusters (Supplementary Fig. 4a). Violin plots of epithelial, stromal, endothelial and immune cell associated genes for those 10 different cell clusters were presented in Supplementary Fig. 4b-e. Clusters which expressed clear markers of non-epithelial cells were removed in the following analyses. Cluster 5 was labeled as endothelial cells based on the expression of Eng, S1pr1 and Emcn (Supplementary Fig. 4d). Cluster 10 was labeled as immune cells based on the expression of Cd74 and Cd72 (Supplementary Fig. 4e). After the removal of above-mentioned non-epithelial cells, we re-ran the Seurat work flow to generate new cell clusters and t-SNE projections (the first 12 principal components were used). A total of 9 clusters were identified (Fig. 5a). We used the FindAllMarkers script in the Seurat package to identify genes that are enriched in a specific cluster (the specific cluster vs. the rest of clusters) with default settings. PCA matrix with the first 12 principal components and clusters from the second Seurat running were taken as input. The cluster representing putative stem cells (Zeb1 expressing cluster 7) was chosen as the root node. Totally, three branches were identified. Heatmaps were plotted by using ComplexHeatmap and ggplot2 package. Similar clusters and the structure of a common origin could also be inferred by using Monocle or Diffusionmap with standard parameters (Supplementary Fig. 5). Codes used are deposited to the website https://github.com/HelenHeZhu/StemCell.

### Human prostate clinical specimens

Freshly dissected human prostate specimens (non-BPH, BPH and prostate cancer specimens) were obtained from the department of Urology at Ren Ji Hospital with informed consent from patients. All Human sample experiments were conducted according to the ethical regulations of Ren Ji Hospital. Human sample collection and handling protocols were approved by the Ren Ji Ethics committee.

### UGM stromal cell preparation

The UGM isolation procedures has been described previously^43^. Briefly, E17 embryos from pregnant SD rats were sacrificed and urogenital sinuses were collected. The UGM was separated from the urogenital sinus epithelium and then digested with 1X collagenase/hyaluronidase solution on a shaker for 90 min at 37℃. The UGM was washed with PBS once and then placed in 0.25%Trypsin/EDTA for 6min at 37℃. UGM single cell suspension was seeded to cell culture dishes and cultured in DMEM supplemented with 10% fetal bovine serum (FBS), 2mM glutamine, 100U/ml penicillin and 100mg/ml streptomycin in vitro for at least 1 week.

### Serial isolation and renal capsule transplantation

We used a previously described procedure to isolate the primary prostate epithelial cells and to perform renal capsule transplantation ^43^. For the first renal capsule transplantation, single or indicated number of Lineage^−^Sca-1^+^CD49f^hi^tdTomato^+^ or Lineage^−^Sca-1^+^CD49f^hi^tdTomato^−^ prostate basal cell was sorted into each well of a 96-well plate via FACS and then examined under a light microscope. The viable sorted cell was mixed with purified rat UGM cells (250,000 cells per graft) in rat tail Collage Type I (4.42mg/ml, Corning, 354236). Next, the mixture was plated as a drop of 10µl into the center of a well of a 6-well plate. The plate was placed in a 37℃ cell culture incubator for at least 60 min to make sure that the Collage type I solidified. Then pre-warmed DMEM (containing 10%FBS) medium was gently pipetted into each well. Each collagen graft was embedded underneath the renal capsule of a nude mouse the next day, For the secondary renal capsule transplantation, primary grafts were digested with 1X collagenase/hyaluronidase solution for 90 min and 0.25% Trypsin/EDTA for 6 min at 37℃. Lineage^−^Epcam^+^Zeb1^+^ or Lineage^−^ Epcam^+^Zeb1^−^ cells were sorted via FACS and then mixed with UGM stromal cells. All serial transplantation grafts were harvested at 8 weeks after implantation. UGS from Zeb1^−/−^ and Zeb1^+/+^ embryos were implanted under the renal capsule of male nude mice. The transplants were collected at 20 days post implantation and sectioned for further analysis.

### Cell lineage tracing

To trace the fate of Zeb1^+^ basal cells, *Zeb1-CreERT2/tdTomato* mice were administrated with 1.25 mg tamoxifen (40mg/mL solution, dissolved into corn oil, Sigma) via intraperitoneal injection at postnatal day 3. Two days and 12 days after the induction, prostates were collected and sectioned for immunofluorescence staining of different prostate epithelial cell markers.

### Organoid culture

We utilized a previously reported protocol to culture organoids *in vitro* and perform organoid frozen sections for immunostaining analysis ^10^. FACS purified Zeb1^+^Basal (Lineage^−^Sca-1^+^CD49f^hi^ tdTomato^+^) and Zeb1^−^Basal (Lineage^−^Sca-1^+^CD49f^hi^ tdTomato^−^) were centrifuged at 350g for 5 min at 4℃ and resuspended in complete organoid culture media with the density of 5000 to 10,000 prostate epithelial cells per 100μL media. The medium was refreshed every 3 days. After approximately 2 weeks, organoids were washed with cold PBS buffer repeatedly to remove the matrigel, collected via centrifugation at 350g for 5min, then dissociated using 0.25% Trypsin/EDTA for 7 min at 37℃. The single cells were centrifuged and resuspended in medium following the aforehand described procedure for a serial passaging. For section preparation, organoids were collected, fixed with 4% PFA for 30min, washed with cold PBS buffer twice, then centrifugated and resuspended with 50μL collagen type I and incubated at 37℃ for 1 hour to solidify the organoid/collagen slurry. The slurry was transferred into 4%PFA solution for fixation, 30% sucrose for dehydration and finally O.C.T for embedding and section. Five-micrometer frozen sections were used for following immunostaining.

### Generation of CRISPR/Cas9 mediated Zeb1 knockout prostate organoids

The lentiCRISPRv2-mCherry plasmid containing cas9 and Cherry fragments was purchased from Addgene (#99154, kindly deposited by Dr.Agata Smogorzewska). Then the Cherry fragment was exchanged for GFP fragment via enzymatic digestion and connection followed by sgRNA cloning. The single guide RNA sequences are ACTGCTTATATGTGAGCTAT in the upstream and GGAACAACCTGAAGTTGACT in the downstream of the sixth exon of the mouse *Zeb1* gene. The Zeb1-cas9 lentivirus was transfected into primary prostate cells using a spinoculation method at 3,000rpm and room temperature for 2 hours. Cells were then transferred to a 15 mL tube and spinned down at 400g for 5min. The pellet was resuspended with the complete organoid culture media and seeded at a density of 10,000 cells per well of a 96-well plate. After 3 weeks of culture, the organoids were harvested and dissociated with pre-warmed 0.25% Trypsin-EDTA. Single cell suspension was stained with anti-EpCAM antibody. EpCAM^+^GFP^+^ cells (sgRNA successfully transfected prostate epithelial cells) were FACS sorted for organoid culture. Organoids were collected when reaching 200μm in diameter, and then fixed, dehydrated and embedded with O.C.T for frozen section preparation.

### RNA extraction and quantitative-PCR analysis

Total RNA was extracted from FACS purified Zeb1^+^ (Lineage^−^Sca-1^+^CD49f^hi^ tdTomato^+^) or Zeb1^−^ basal cells (Lineage^−^Sca-1^+^CD49f^hi^tdTomato^−^) using a RNeasy micro kit (Qiagen). cDNA was synthesized using the PrimeScript RT Reagent Kit (Takara). qPCR was conducted using SYBR Premix Ex Taq (Takara). Relative transcript abundance was quantified by the comparative CT method using Actin as an internal reference gene. Primers utilized in RT-PCR were shown in Supplementary Table 4.

### Immunofluorescence staining

Human or mouse prostates were fixed in 4% paraformaldehyde for 20 min and dehydrated overnight in 30% sucrose solution. Tissues were embedded in Optional Cutting Temperature (O.C.T.) compound and placed in a −80℃ refrigerator for 10 minutes or longer depending on the tissue size. Frozen sections were cut at a thickness of 6 um. Sections were washed with PBS buffer and placed into 0.01M sodium citrate (PH 6.0) buffer for heat-induced antigen retrieval. Slides were then subjected to a blocking step in 10% donkey serum solution for 1 hr at room temperature. Primary antibodies, diluted in 1% donkey serum, were added to sections overnight at 4℃. After thorough wash with PBS buffer, secondary antibodies were applied and incubated for 1 hr at room temperature. Sections were rinsed at least three times with PBS before being mounted by Vector Shield mounting medium containing DAPI. Antibodies used in the study is listed in Supplementary Table 5.

### Immunohistochemistry

Paraffin-embedded prostate tissue sections were deparaffinized and rehydrated following conventional methods. Harris modified hematoxylin solution (Sigma) was applied to sections for 5 min followed by water washing. Then 1% Eosin solution (Sigma) was applied to sections for 3 min. After thorough wash, sections were mounted using neutral balsam.

### Castration and androgen replacement

Zeb1 reporter mice were surgically castrated. Three weeks later, mice were given dihydrotestosterone (MCE, HY-A0120) dissolved in sterile corn oil via intraperitoneal injection twice a day (50ug/d) for prostate regeneration. Prostate collected at indicated regeneration stages (regressed, regenerated, and recovered stages post testosterone administration) for analysis.

### Statistical analysis

We used the ImageJ 1.46r software to quantify positive stained cells. All statistical analysis was evaluated using a two-tailed Student’s T-test and a p-value<0.05 was defined to be significant.

### Data and materials availability

The single cell RNA-seq raw data used for this study are available at the GEO web with the accession number GSE111429. Codes used are deposited to the website https://github.com/HelenHeZhu/StemCell. All other data is available in the main text or the supplementary materials.

