## Supplemental info.(including supplemental figures and figure legends) for "Identification of a basal stem cell subpopulation in the prostate via functional, lineage tracing and single-cell RNA-seq analyses"

**Contents:**

Supplementary Fig.1

Supplementary Fig.2

Supplementary Fig.3

Supplementary Fig.4

Supplementary Fig.5

Supplementary Fig.6

Supplementary Table 1

Supplementary Table 2

Supplementary Table 3

Supplementary Table 4

Supplementary Table 5

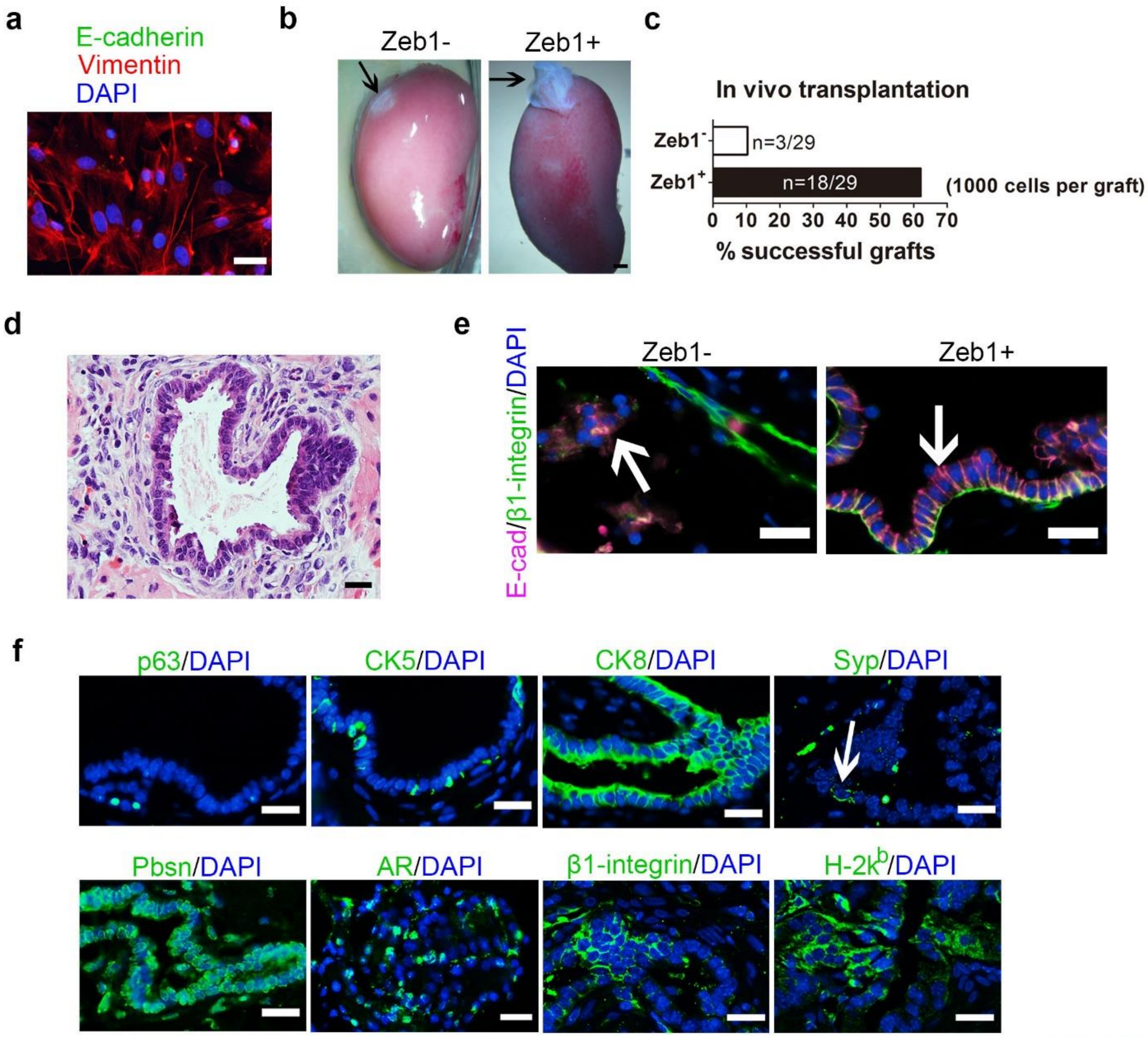
a
E-cadherin
Vimentin
DAPI
b
Zeb1-
Zeb1+
c
In vivo transplantation
Zeb1⁻
n=3/29
Zeb1⁺
n=18/29
(1000 cells per graft)
0 10 20 30 40 50 60 70
% successful grafts
d
e
E-cad/β1-integrin/DAPI
Zeb1-
Zeb1+
f
p63/DAPI
CK5/DAPI
CK8/DAPI
Syp/DAPI
Pbsn/DAPI
AR/DAPI
β1-integrin/DAPI
H-2k^b/DAPI

**Figure S1**

**Supplementary Fig. 1| Zeb1$^{+}$ basal cells are enriched for multipotent prostate basal stem cells**

**a,** Immunostaining of E-cadherin and Vimentin to validate the purity of UGM cells separated from UGS.

**b,** Prostate tissue generated from Lineage$^{-}$Sca-1$^{+}$CD49f$^{hi}$ Zeb1$^{+}$ prostate cells at 2 months after renal capsule implantation. (implanted cell number: 1000)

**c,** Quantification of prostate tissue generation incidence from 1000-cell transplants.

(All scale bars=20 μm.)

**d,** H&E staining of 1000-cell implants shows well differentiated prostate epithelial tubules.

**e,** Immunostaining for β1-integrin (green) and E-cadherin (red) on sections of renal capsule implants displays viable mouse specific prostate epithelial cells in Zeb1$^{-}$ grafts.

**f,** Immunostaining of p63, CK5, CK8, Syp, Pbsn, AR, mouse-specific β1-integrin and C57BL/6 donor-specific H-2k$^{b}$ on sections of 1000-cell implants. (All scale bars=20 μm.)

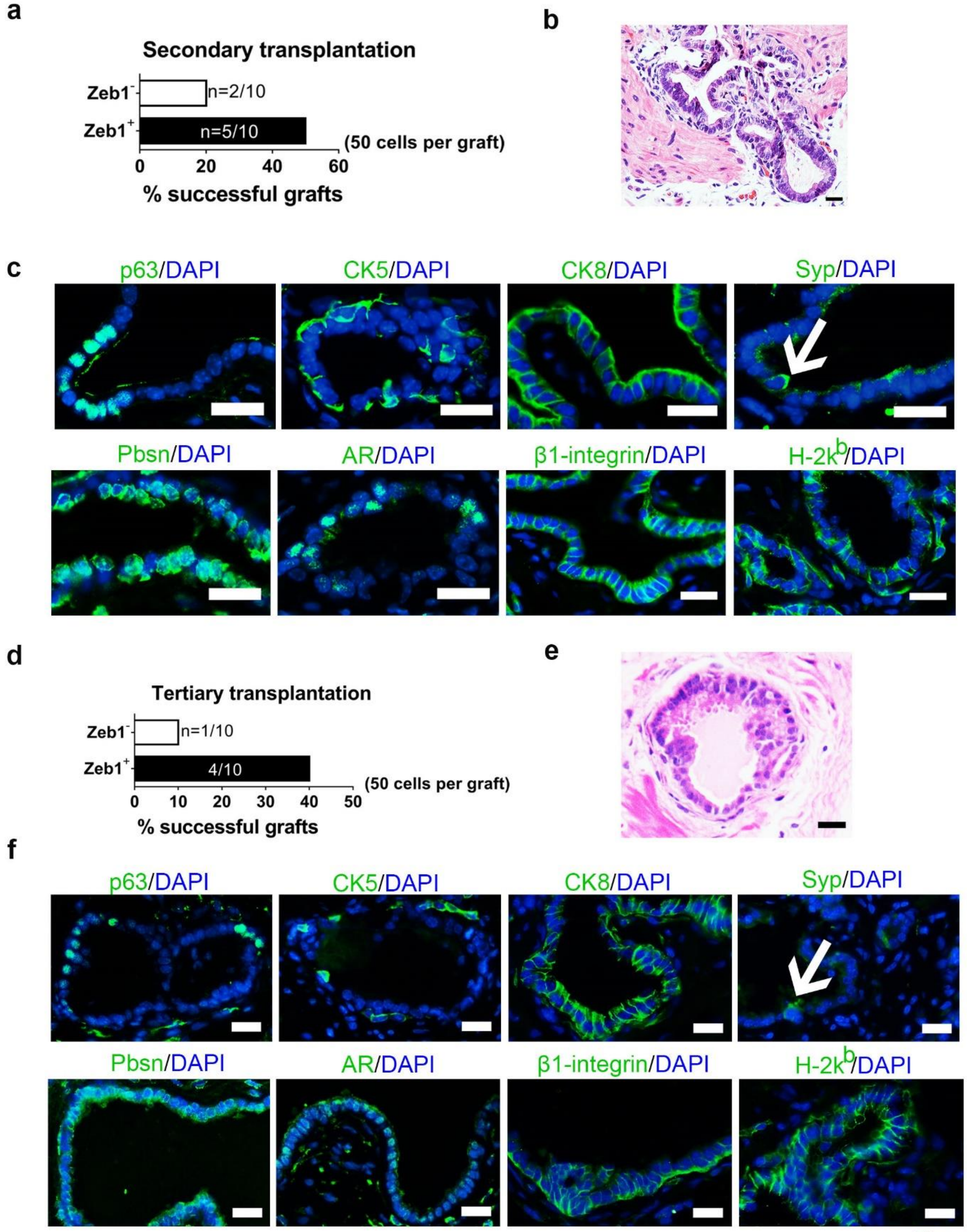
a
Secondary transplantation
Zeb1-
n=2/10
Zeb1+
n=5/10
(50 cells per graft)
0
20
40
60
% successful grafts
b
c
p63/DAPI
CK5/DAPI
CK8/DAPI
Syp/DAPI
Pbsn/DAPI
AR/DAPI
β1-integrin/DAPI
H-2kb/DAPI
d
Tertiary transplantation
Zeb1-
n=1/10
Zeb1+
4/10
(50 cells per graft)
0
10
20
30
40
50
% successful grafts
e
f
p63/DAPI
CK5/DAPI
CK8/DAPI
Syp/DAPI
Pbsn/DAPI
AR/DAPI
β1-integrin/DAPI
H-2kb/DAPI

Figure S2

**Supplementary Fig. 2| Zeb1$^+$ basal cells can self-renew and generate functional prostates *in vivo* in a serial renal capsule transplantation assay.**

**a,** Zeb1$^+$EpCAM$^+$ cells harvested from first generation of renal capsule implantation were able to form prostates in secondary transplantations. (50 prostate cells per graft)

**b,** H&E staining of secondary transplants derived from Zeb1$^+$ prostate epithelial cells showing differentiated prostate epithelial tubules.

**c,** Immunostaining of p63, CK5, CK8, Syp, Pbsn, AR, mouse-specific β1-integrin and C57BL/6 donor-specific H-2k$^b$ on sections of secondary transplants derived from Zeb1$^+$ prostate epithelial cells. (Syp, indicated by an arrow, All scale bars=20μm.)

**d,** Zeb1$^+$EpCAM$^+$ cells harvested from first generation of renal capsule implantation were able to form prostates in tertiary transplantations. (50 prostate cells per graft)

**e,** H&E staining of tertiary transplants derived from Zeb1$^+$ prostate epithelial cells showing differentiated prostate epithelial tubules.

**f,** Immunostaining of p63, CK5, CK8, Syp, Pbsn, AR, mouse-specific β1-integrin and C57BL/6 donor-specific H-2k$^b$ on sections of tertiary transplants derived from Zeb1$^+$ prostate epithelial cells. (Syp, indicated by an arrow, All scale bars=20μm.)

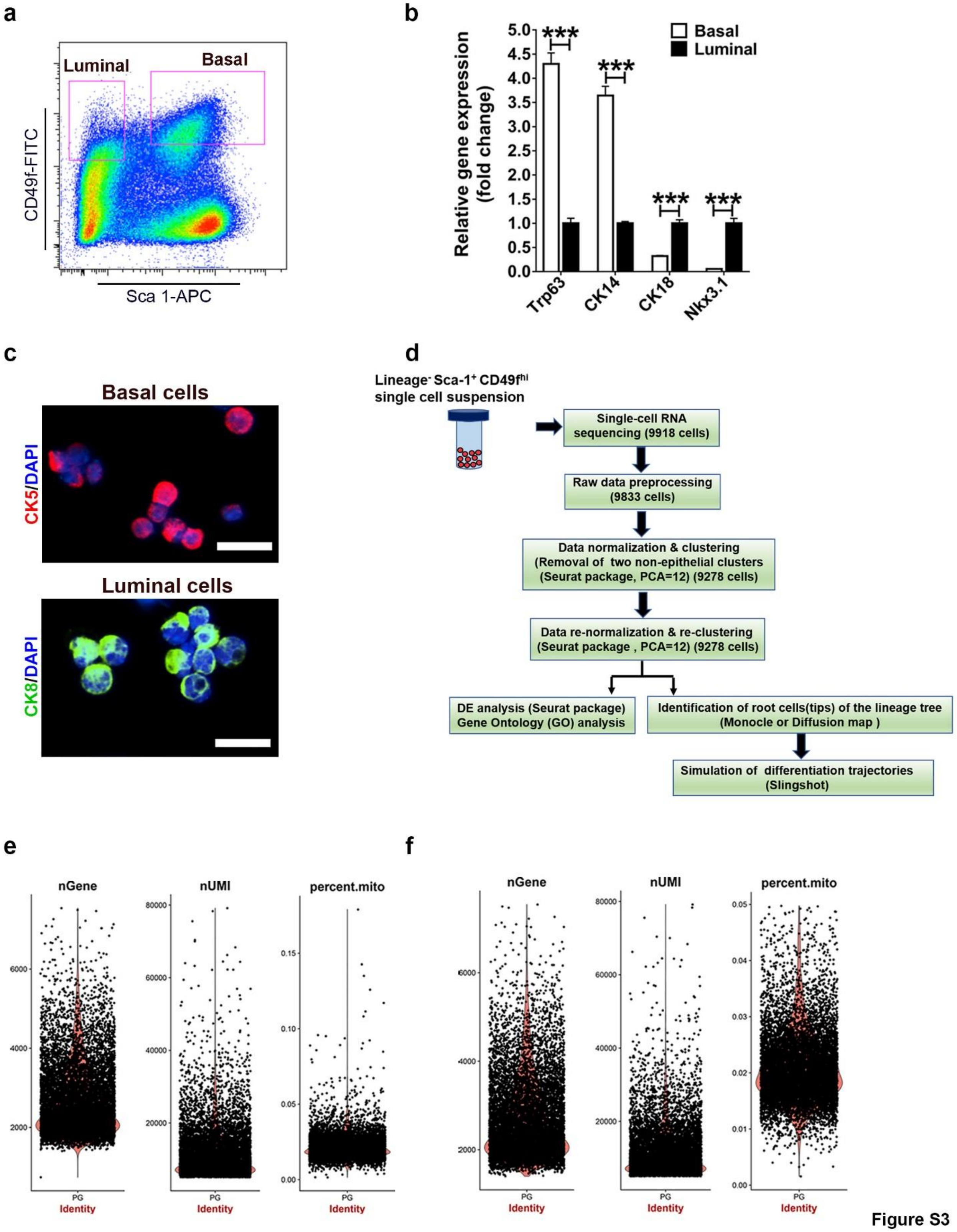
a
Luminal
Basal
CD49f-FITC
Sca 1-APC
b
Relative gene expression (fold change)
Basal
Luminal
Trp63
CK14
CK18
Nkx3.1
c
Basal cells
CK5/DAPI
Luminal cells
CK8/DAPI
d
Lineage⁻ Sca-1⁺ CD49f^hi single cell suspension
Single-cell RNA sequencing (9918 cells)
Raw data preprocessing (9833 cells)
Data normalization & clustering (Removal of two non-epithelial clusters (Seurat package, PCA=12) (9278 cells)
Data re-normalization & re-clustering (Seurat package , PCA=12) (9278 cells)
DE analysis (Seurat package) Gene Ontology (GO) analysis
Identification of root cells(tips) of the lineage tree (Monocle or Diffusion map )
Simulation of differentiation trajectories (Slingshot)
e
nGene
nUMI
percent.mito
Identity
f
nGene
nUMI
percent.mito
Identity

**Figure S3**

**Supplementary Fig. 3| Single cell RNA-seq data pre-processing**

**a,** FACS plots displaying the gating of basal ($Lineage^{-}Sca\text{-}1^{+}CD49f^{hi}$) or luminal ($Lineage^{-}Sca\text{-}1^{-}CD49f^{lo}$) prostate epithelial cells.

**b,** qRT-PCR quantification of basal (Trp63, CK14) and luminal (CK18, Nkx3.1) cell markers in sorted basal and luminal cells. RNA expression levels were normalized to luminal cells (n=3).

**c,** Immunostaining results show CK5 expression in sorted basal cells and CK8 expression in sorted luminal cells.

**d,** The flow chart for single-cell RNA sequencing data analysis.

**e, f,** Violin plots displaying the distribution of detected numbers of gene, unique molecular identifiers (UMI) and percentage of UMIs mapped to mitochondrial genes before (**e**) and after (**f**) removal of low quality cells (cells with less than 5000 UMIs or less than 1400 genes detected or more than 5% UMI mapped to mitochondria genes were removed).

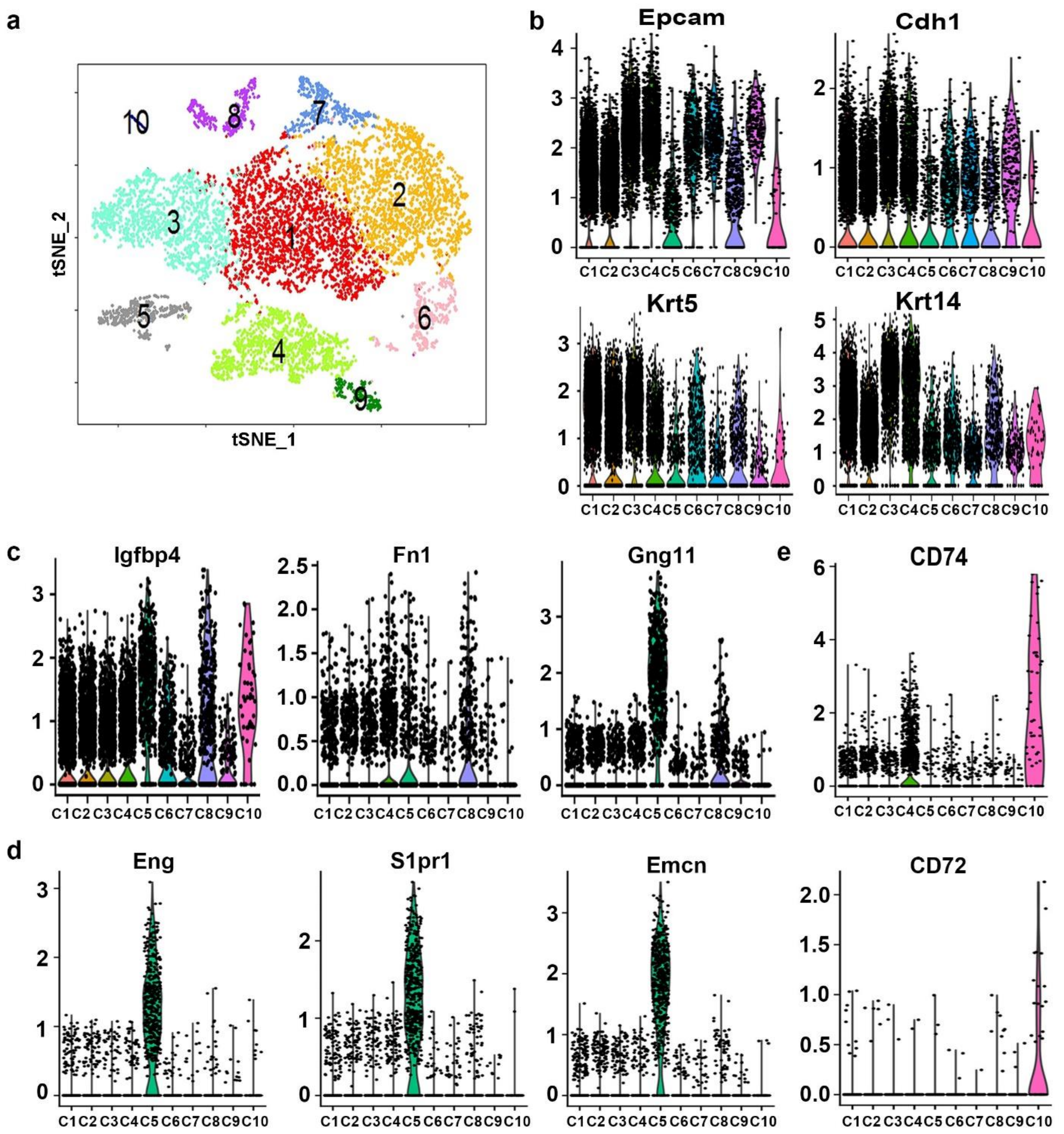
a
tSNE_2
tSNE_1
b
Epcam
Cdh1
Krt5
Krt14
c
Igfbp4
Fn1
Gng11
e
CD74
d
Eng
S1pr1
Emcn
CD72
C1 C2 C3 C4 C5 C6 C7 C8 C9 C10

Figure S4

**Supplementary Fig. 4| Single cell RNA-seq data pre-processing followed by clustering.**

**a,** A Seurat package and the first 12 principal components were applied to generate 10 different and stable clusters based on differential expressing genes among 9278 mouse prostate basal cells.

**b-e,** Violin plots showing expression difference for epithelial (**b**), stromal (**c**), endothelial (**d**) and immune cell (**e**) related genes for the 10 cell clusters.

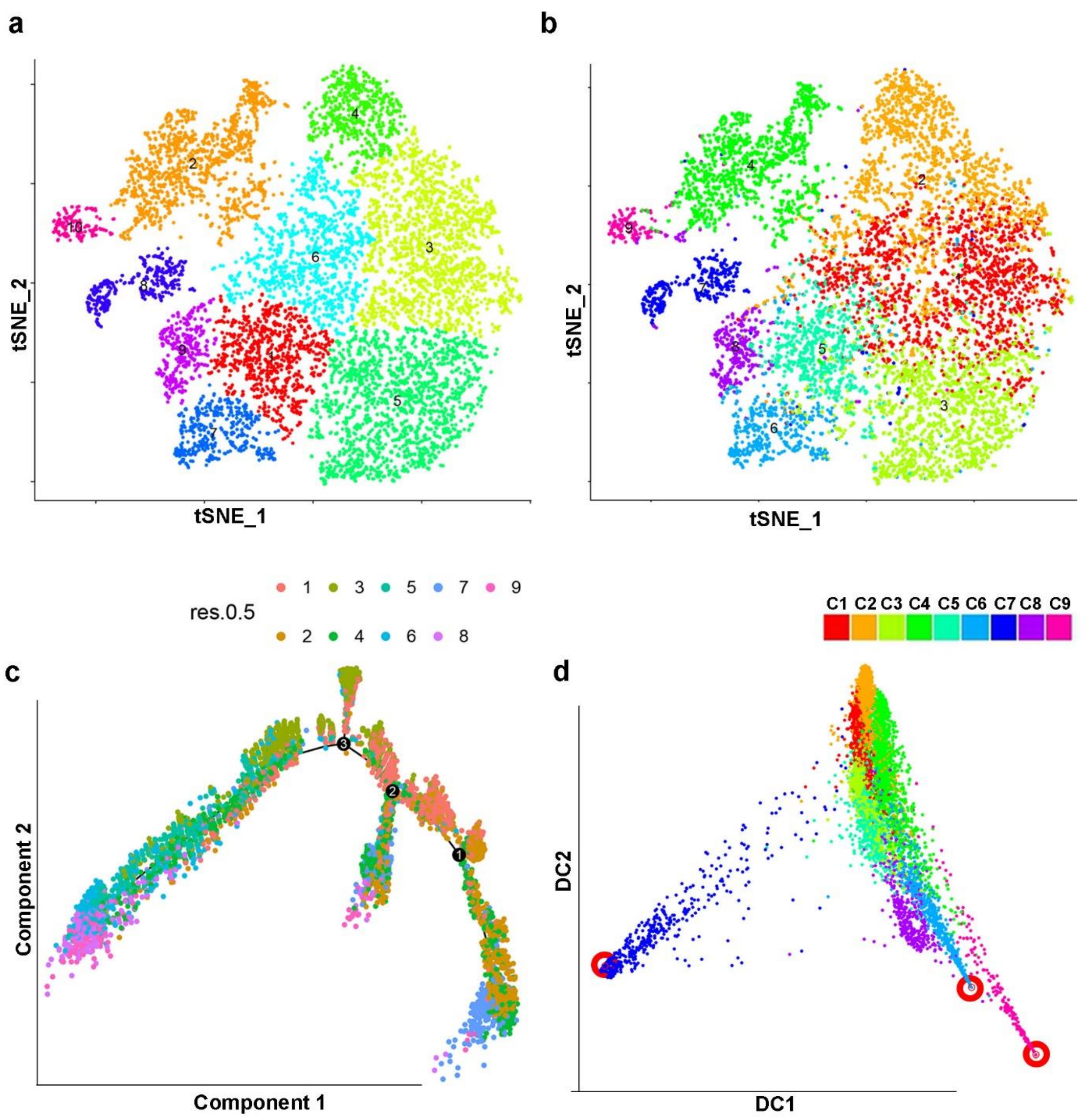
a
b
tSNE_2
tSNE_1
res.0.5
1 3 5 7 9
2 4 6 8
C1 C2 C3 C4 C5 C6 C7 C8 C9
c
d
Component 2
Component 1
DC2
DC1

Figure S5

**Supplementary Fig. 5| Single cell RNA-seq data pre-processing followed by simulation of lineage tree using monocle and diffusion map packages.**

**a, b,** t-SNE plots displaying the cell clusters inferred by Monocle (**a,** the ten cell clusters) and Seurat (**b,** the nine cell clusters). Compared to Seurat results, monocle similarly separated the C3, C4, C5, C6, C7, C8, and C9, but had some differences in the C1 and C2 separation. C7 from Monocle (**a**) or Seurat (**b**) analyses is the EMT-like cell cluster.

**c,** Differentiation trajectory was simulated by Monocle colored by cluster assignment (Colors were corresponding to the cluster IDs which were generated by Seurat). The data showed that C7 is at the tip position of one branch (labeled in skyblue at the low right).

**d,** Cell dimension reduced by DiffusionMap (different colors represented clusters defined by Seurat). The red hollow circle showed the three tips identified by DiffusionMap under default parameters.

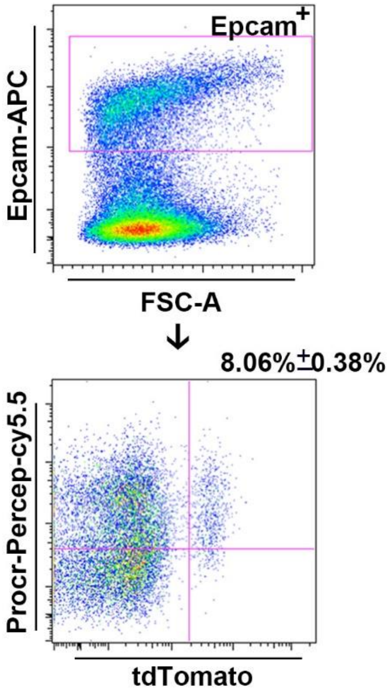


Figure S6

**Supplementary Fig. 6| Zeb1$^{+}$ epithelial cell is a subpopulation of Procr expressing prostate epithelial cells.**

FACS analysis of the expression of Zeb1/tdTomato and Procr in EpCAM$^{+}$ prostate epithelial cells from Zeb1/tdTomato reporter mice. (n=4)
